# happi: a Hierarchical Approach to Pangenomics Inference

**DOI:** 10.1101/2022.04.26.489591

**Authors:** Pauline Trinh, David S. Clausen, Amy D. Willis

**Affiliations:** Department of Environmental & Occupational Health Sciences, University of Washington; Department of Biostatistics, University of Washington

**Keywords:** shotgun metagenomics, metagenome-assembled genomes, microbiome, statistical models, hypothesis testing

## Abstract

Recovering metagenome-assembled genomes (MAGs) from shotgun sequencing data is an increasingly common task in microbiome studies, as MAGs provide deeper insight into the functional potential of both culturable and non-culturable microorganisms. However, metagenome-assembled genomes vary in quality, and may contain omissions and contamination. These errors present challenges for detecting genes and comparing gene enrichment across sample types. To address this, we propose `happi`, an approach to testing hypotheses about gene enrichment that accounts for genome quality. We illustrate the advantages of `happi` over existing approaches using published Saccharibacteria MAGs and via simulation.

## 1 Background

Members of the same bacterial species can display a wide variety of different phenotypes, and intra-species variation in pathogenicity, virulence, drug resistance, environmental range, and stress response has been observed across the tree of life [16, 19, 23, 30, 34]. Variation in phenotypes can in part be explained by genotypic variation, which is also considerable because mechanisms of genetic recombination in bacteria facilitate large genetic variation even within narrow organismal groups. For example, of 7,385 gene clusters observed in a study of 31 genomes in the genus *Prochlorococcus*, only 766 gene clusters were detected in all genomes [8]. We refer to the set of genes shared by all members of a clade as the *core genome* and we refer to the set of genes not shared by all members as the *accessory genome* [33]. Together, these sets of genes comprise a clade’s *pangenome*: the entire collection of genes present in one or more organisms within the clade. In this paper, we describe a novel tool for pangenome analysis. Our tool is a statistical method to model the association between gene presence and covariates (predictors). Our method offers interpretable parameter estimates, a fast algorithm for estimation, and a flexible hypothesis testing procedure.

While cultivation-based studies have historically been used to study the gene content of bacteria, it has become increasingly common to employ shotgun metagenomics to study bacterial genomes and communities. Shotgun metagenomic sequencing involves untargeted sequencing of all DNA in an environment, enabling the study of genomes in their environmental context. Short reads from shotgun sequencing can be assembled into contigs and binned into metagenome-assembled genomes (MAGs), which represent a partial reconstruction of an individual bacterial genome. Despite major advances in methods for binning MAGs, MAGs can contain two types of errors. First, there can be genes that are truly present in the genome the MAG represents, but are unobserved in a MAG. Common reasons for this error include inadequate sequencing depth, high diversity in the metagenomes under study, and the inherent limitations of short read sequencing for reconstructing repetitive regions [10, 22, 24, 29, 37]. A second type of error in MAGs is erroneously observed genes: genes that are included in a MAG that are not truly present in the originating genome. This phenomenon is often referred to as contamination. The use of automated binning tools in the absence of manual inspection and refinement can lead to elevated rates of contamination. For example, the identification of contaminating contigs from manual refinement of MAGs produced by a massive unsupervised genome reconstruction effort removed 30 putative functions from a single contaminated genome [5, 20].

To address the challenges that contaminating and unobserved genes create for detecting enriched genes, our proposed method incorporates information about each genome’s quality. Under our proposed model, a gene may be unobserved in a genome either because the gene is not present in the source genome, or because it could not be recovered from the obtained sequencing data. If, for example, the coverage of short reads across the genome was high and most of the expected core genes were observed, then the lack of detection of a given gene is more likely attributable to its true absence. The user can select which variables they believe to be the most informative for genome quality in their dataset. We develop estimators of the parameters of our model, discuss interpretation of model parameters, propose a hypothesis testing approach, and illustrate the performance of our model on shotgun sequencing and simulated data.

## 2 Results

### 2.1 A Hierarchical Model for Gene Presence

We present a hierarchical model for the association between bacterial gene presence and covariates of interest (e.g., host treatment status, environment of origin, relevant confounders, etc.). We consider observations on *n* genomes, which could be either metagenome-assembled genomes, isolate genomes, reference genomes, or any combination. Let *Y_i_* be an indicator variable for the gene of interest being *observed* in genome *i*, *Y_i_* = 1 if the gene is observed in genome *i* and *Y_i_* = 0 otherwise. However, we are not interested in whether the gene is *observed* in each genome – we are interested in whether it is *present* in each genome. To this end, we define λ_*i*_ to be a latent (unobserved) random variable that indicates if the gene is truly present in genome *i* (λ_*i*_ = 1 if present).

We propose a logistic model to connect gene presence to covariate vector 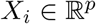:

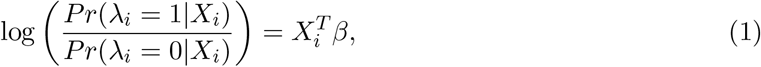

where the λ_*i*_’s are conditionally independent given *X_i_* and follow a Bernoulli distribution. Therefore, when comparing groups of genomes that differ by one unit in *X._k_* but are alike with respect to *X*._1_, *X*._2_,…, *X*._,*k*–1_, *X*._,*k*+1_,…, *X*._*p*_, *β_k_* gives the difference in the log-odds that the gene will be present between these two groups of genomes. To connect λ_*i*_ to *Y_i_* we propose the following model

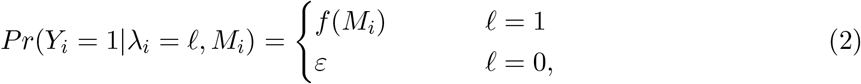

where *Y_i_* are conditionally independent Bernoulli distribution random variables; *ε* is the probability that a gene is observed in a genome in which it is absent (e.g., due to contamination or crosstalk); 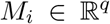 is a vector of genome quality covariates; and 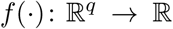 is a flexible function to connect quality variables to the probability of detecting a present gene. Relevant quality variables are context-dependent and could include coverage of the gene from metagenomic read recruitment, completion (percentage of single copy core genes observed in the genome), redundancy (percentage of single copy core genes observed more than once in the genome), and an indicator for the genome originating from an isolated bacterial population.

### 2.2 Parameter Estimation

The latent variable structure of our model makes the Expectation-Maximization Algorithm [9] an appealing choice for estimating unknown parameters *θ* = (*β*, *f*). Because we do not observe 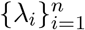, *ε* and *f* are not, in general, jointly identifiable. Therefore, we treat *ε* as a hyperparameter that can be fixed by the user or leveraged for sensitivity analyses. To improve stability of parameter estimates, we impose a Firth-type penalty on *β*. The complete data penalized log-likelihood is linear in λ_*i*_, which allows us to simplify the expected complete data penalized log-likelihood at step *t* of an EM iteration as

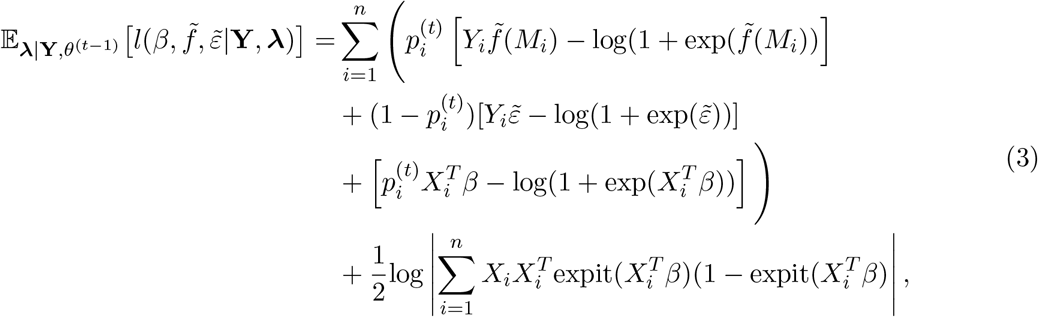

where 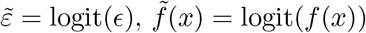 for all *x*, and 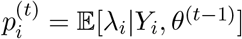 can be simplified as

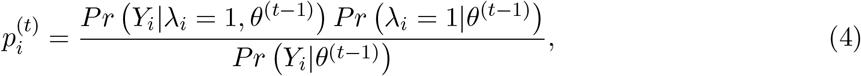

where the terms in the numerator are given in (1) and (2), and the denominator is given by

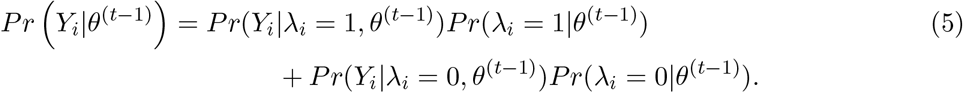

We maximize the expected complete data penalized log-likelihood separately for *β* and *f*. Owing to the form of the expected complete data penalized log-likelihood, efficient algorithms exist to perform each of these maximizations. Optimizing (3) with respect to *β* is equivalent to fitting a binomial generalized linear model with logit link function for outcomes 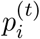 via Firth-penalized maximum likelihood, and we find Newton’s method to be stable and fast for this purpose.

Optimizing for *f* depends on the class of functions in which *f* falls. We investigated two flexible non-parametric options for 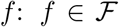, where 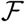 is the class of bounded non-decreasing functions that map from 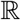 to 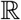, and 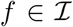 where 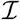 is the class of linear combinations of *k* I-spline basis functions and a constant function where all basis functions have nonnegative coefficients. Both 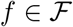 and 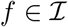 result in a monotone estimate for *f*. To obtain the EM update for 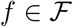, we use the primal active set algorithm of `isotone` [7] with custom loss function given by the first term in (3) plus a penalty term 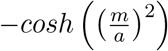 to prevent 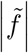 from growing without bound. We found that setting *a* = 50 gives a sensible tradeoff between algorithm convergence and numerical stability. To obtain the EM update for 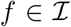, we fit a logistic regression on 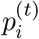 with predictors consisting of an I-spline basis with all non-intercept coefficients constrained to be nonnegative. We use the I-spline basis functions implemented in `splines2` [35]. In an analysis where we used short-read subsampling to approximate an empirical *f*, we found that 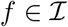 outperformed 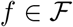 (see Section 4.2.2), and for that reason we consider 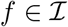 throughout the remainder of this manuscript. We run the estimation algorithm for *t*_max_ steps or until the relative increase in the log-likelihood is below threshold Δ for 5 consecutive steps.

### 2.3 Hypothesis Testing

To enable inference on the odds that a gene will be present in groups of genomes that differ in their covariate attributes, we construct a hypothesis test for null hypotheses of the form **A***β* = *c* for 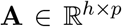 and 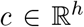 where rank(**A**) = *h*. This allows testing of null hypotheses including *β_k_* = 0 (the odds that the gene will be present are equal when comparing groups of genomes that differ in *X*._k_ but are alike with respect to *X*._1_, *X*._2_,…, *X*._,*k*–1_, *X*._;*k*+1_,…, *X*._,*p*_). We propose to use a likelihood ratio test for **A***β* = *c*, rejecting *H*_0_ at level *α* if 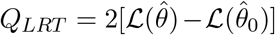 exceeds the upper 100*α*% quantile of a 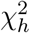 distribution, where 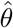 is the maximum likelihood estimate of *θ*; 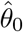 is the maximum likelihood estimate of *θ* under the null hypothesis; and 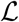 is the log-likelihood function.

### 2.4 Data Analysis: Saccharibacteria MAGs

We consider a publicly-available dataset of *n* = 43 non-redundant Saccharibacteria (TM7) MAGs recovered from supragingival plaque (*n* = 27) and tongue dorsum (*n* = 16) samples of seven individuals from [28] (see Section 4 for more information). The wide variation in mean coverage across the MAGs (1.07 – 26.35×) makes this an appealing dataset on which to illustrate our quality variable-adjusting pangenomics method.

We consider methods that allow us to test the null hypothesis that the probability (equivalently, odds) that a gene is present in Saccharibacteria genomes are equal for tongue and plaque-associated genomes. The alternative hypothesis is that the probabilities differ. We compare our proposed method (`happi`: a Hierarchical Approach to Pangenomics Inference) with three competitors: a logistic regression model for *Y_i_* with a likelihood ratio test (GLM-LRT); a logistic regression model for *Y_i_* with a Rao test (GLM-Rao); and Fisher’s exact test (Fisher). Note that these latter three methods test hypotheses about the odds that a gene is observed, while our proposed approach tests hypotheses about the odds that a gene is present, but we believe that results can be reasonably compared between these methods. We consider a single quality variable *M_i_* for our analysis with `happi`: mean coverage across genome *i*. Our primary comparison is with GLM-Rao, which is the method currently implemented for pangenomics hypothesis testing in anvi’o [28]. We also note that the results from GLM-Rao and GLM-LRT are highly correlated, especially for larger p-values.

Different methods identified different differentially present genes. Out of 713 COG functions tested, `happi` identified 171 differentially present genes when controlling false discovery rate at the 5% level; GLM-LRT identified 219 genes; GLM-Rao identified 175 genes; and Fisher identified 146 genes. Our proposed method calculated lower p-values for 20%, 35% and 85% of genera compared to GLM-LRT, GLM-Rao, and Fisher’s test. We show results from 6 specific model estimates in Figure 1: 3 genes for which `happi` produced greater p-values than GLM-Rao (upper panels), and 3 genes for which it produced smaller p-values than GLM-Rao (lower panels). In all instances where `happi` produced greater p-values than GLM-Rao, non-detections generally occurred in genomes with low mean coverage. GLM-Rao does not account for coverage information, and so unlike `happi`, it can conflate gene absence with non-detections due to quality. We believe that statements about significance should be moderated when detection patterns can be attributable to quality variables, and therefore that it is reasonable that p-values are larger in these three cases. In contrast, `happi` produced smaller p-values than GLM-Rao in instances when non-detections occurred for greater coverage MAGs, or broadly across the range of MAG coverage (lower panels). In these instances, differences in detection are less likely to be attributable to quality factors, and it is reasonable that the significance of findings can be strengthened by including data on quality variables.

**Figure 1:**
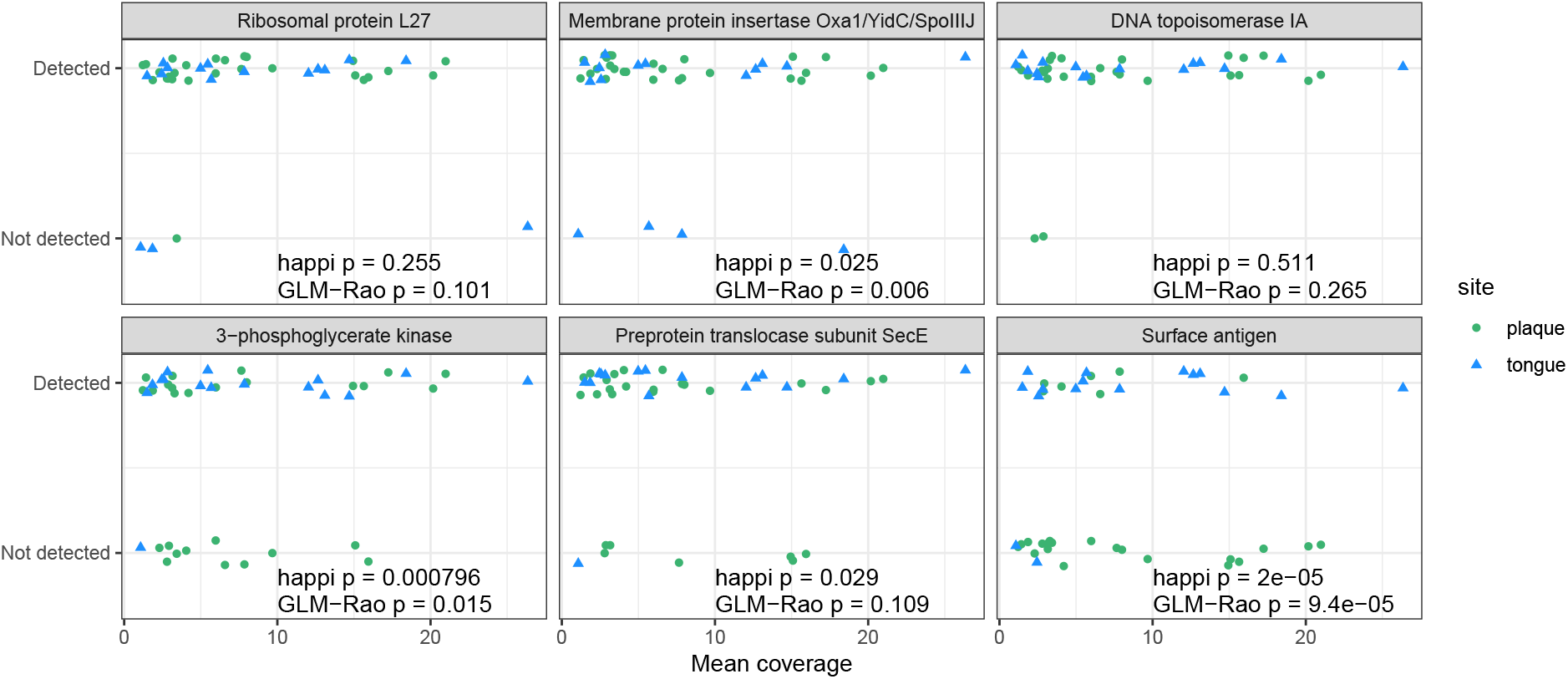
We test the null hypothesis that the probability that a gene is present are equal for tongue and plaque-associated Saccharibacteria genomes. The top 3 panels show genes for which our proposed method resulted in greater p-values than existing methods, and the lower 3 panels shows genes for which our proposed method resulted in smaller p-values than existing methods. Our method reduced p-values when differences in detection cannot be attributed to genome quality factors (here, coverage), and increased p-values in situations when non-detection may be conflated with lower quality genomes. Points have been jittered vertically to separate observations.

### 2.5 Simulation Study

Finally, we investigate the performance of our approach by evaluating its Type 1 error rate and power. To generate data that most realistically reflects the relationship between coverage and gene detection in shotgun metagenomics studies, we construct *f*(·) for use in this simulation by subsampling short-reads from host-associated *E. coli* genomes ([1]; see Section 4.2 and Figure 3). We consider *q* = 1 and *q* = 2, and let 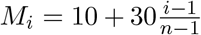, *X*_*i*1_ = 1, 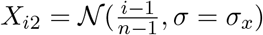 and *ϵ* = 0. *σ_x_* is a parameter that controls the degree of correlation between *M_i_* and *X*_*i*2_, with larger values resulting in less correlation between quality variables and the predictor of interest. We simulate data according to the model described in (1) and (2), with *β* = (0, 0)^*T*^ for Type 1 error simulations and *β* = (0, *β*_1_)^*T*^ with *β*_1_ ≠ 0 for power simulations. Note that because *X*_*i*1_ is continuous, a Fisher’s exact test cannot be applied in this setting.

The results of Type 1 error rate simulations are shown in Figure 2 (left panels). We only show results for GLM-Rao because GLM-LRT and GLM-Rao produced highly similar p-values (mean squared difference 1.3 × 10^-5^, correlation = 0.99996, *n_sim_* = 3000). Notably, the logistic regression methods are anti-conservative, and do not control Type 1 error rates at nominal levels.

**Figure 2:**
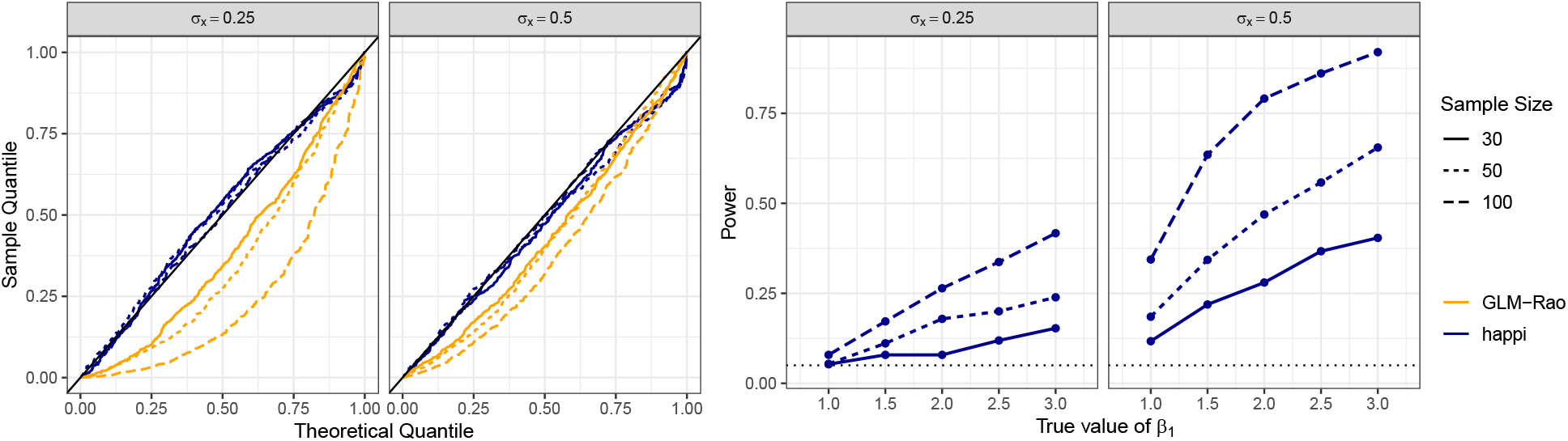
Simulations can be useful for evaluating the Type 1 and Type 2 error rates of methods for testing statistical hypotheses. (left) We find that logistic regression methods do not control Type 1 error, while `happi` behaves control Type 1 error at nominal levels. (right) We evaluate the power of `happi` to reject a false null hypothesis, finding that larger samples have greater power. In situations with greater correlation between quality variables and the covariate if interest, `happi` exhibits comparatively lower power.

**Figure 3:**
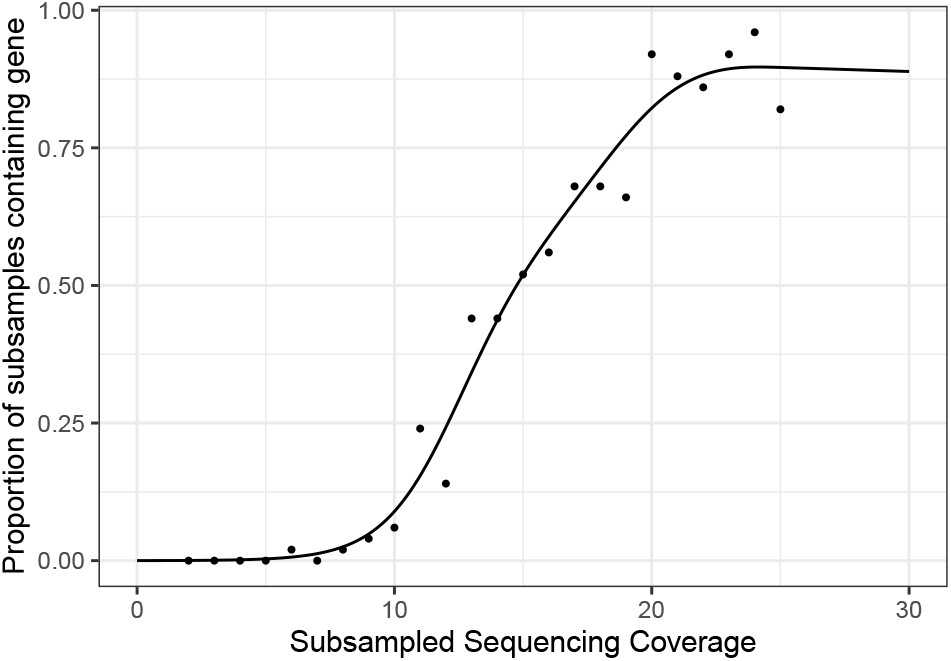
We subsampled reads from a publicly available *E. coli* isolate genome to understand the impact of coverage on the probability of detecting a gene, finding that the probability of detection increases with coverage. We use a nonparametric smoother to interpolate this curve and use it as the true function *f* in our simulations.

For example, for a 5%-level test, Type 1 error rates for GLM-LRT range from 8.6% (*n* = 30 and *σ_x_* = 0.5; 95% CI: 6.1–11.1%) to 31.6% (*n* = 100 and *σ_x_* = 0.25; 95% CI: 27.5–35.7%). Stated differently, under *H*_0_, GLM-LRT will return p-values that are usually too small, leading to more frequent incorrect conclusions of an association. In contrast, `happi` does control the Type 1 error rate, behaving near-exactly. We estimate that `happi`’s Type 1 error rates for a 5% test when *n* = 30 and *σ_x_* = 0.5 is 5.2% (95% CI: 3.3–7.2%), and when *n* = 100 and *σ_x_* = 0.25, `happi`’s empirical Type 1 error rate is 6.0% (95% CI: 3.9–8.1%). Greater correlation between the quality variable (coverage) and the covariate of interest leads to greater anti-conservativeness for logistic regression methods, which incorrectly attribute differences in gene presence to the covariate of interest. However, `happi` appears to control Type 1 error across the range of *σ_x_* investigated here.

We show the power of `happi` to correctly reject a null hypothesis at the 5% level in Figure 2 (right panels). We do not evaluate power for GLM-Rao and GLM-LRT because they have uncontrolled Type 1 error rates, making them invalid tests. We observe that the power of `happi` to reject a false null hypothesis increases with the effect size and sample size, but decreases with greater correlation between *M_i_* and *X*_*i*1_. Stated differently, `happi` has low power to detect true associations between gene presence and covariates of interest when covariates are correlated with genome quality, though this can be remedied with larger sample sizes.

Taken together, these results show that `happi` is robust to potential correlation between covariates of interest and genome quality. This is not the case for logistic regression-based methods, which cannot distinguish between differential gene presence due to genome quality and differential gene presence due to associations with covariates. No method will perform well under the alternative with small sample sizes and high correlation (see Figure 2, third panel), but `happi` has some power for large sample sizes and large effect sizes in this setting, and controls Type 1 error at nominal levels regardless of the sample size.

## 3 Discussion

Many tools exist to study associations between microbial genome variation and microbial or host phenotypes [4, 6, 11, 18, 27]. Studies investigating the association between microbial genomes and phenotypes are often referred to as microbial genome-wide association studies (mGWAS) [21, 25]. Most mGWAS tools have been developed for the analysis of pure microbial isolates, and do not account for differential genome quality in genomes analyzed collectively. mGWAS tools may be better-suited when the hypothesized causal direction is that the presence of genetic features gives rise to a phenotypic characteristic, and not the reverse. In this paper, we propose and validate a novel method (`happi`) to understand how non-microbial variation (e.g., environmental variation) is associated with microbial genome variation. The implied direction of modeling is reversed in our model compared to mGWAS models: our response variable is gene presence rather than phenotype. This allows interrogation of questions about factors influencing selection pressures on genomes, rather than questions about the impact of the microbiome on phenotypic outcomes.

We view the main advantage of `happi` as its use of data about genome quality factors. To support the increasing use of shotgun metagenomic data to recover fragmented microbial genomes, researchers need methods that are capable of analyzing incomplete and imperfect genomes. While we are not aware of methods for modeling gene enrichment in MAGs, we offer comparisons to commonly used methods for analyzing near-complete genomes, such as Fisher’s exact test (used by PanPhlAn3 [2, 26] and Scoary [4]) and logistic regression (used by anvi’o [12, 28]; see also [3]). In situations where differences in gene detection can be attributed to differences in genome quality, `happi` correctly infers that gene enrichment is ambiguous, and correspondingly identifies associations as less significant compared to competitor methods. However, in situations where genome quality cannot explain gene detection patterns, `happi` has greater precision than other methods and produces smaller p-values. We show via simulation that the advantages of `happi` are most pronounced when there is correlation between covariates and quality variables.

Results generated from `happi` are easily interpretable with reasonable run times on a modern laptop without parallelization, averaging 1.04 seconds per gene over 713 genes in *n* = 43 samples with *t*_max_ = 1000 and Δ = 0.01 on a 2.6 GHz i7 processor with 16 GB RAM. Since genes are treated independently, this analysis can be trivially parallelized, and furthermore, accuracy in estimation can be traded off for reduced runtime by reducing *t*_max_ or increasing Δ.

We suggest several avenues for further research. The first is to study the impact of experimental design on the statistical power of our proposed hypothesis testing procedure. Researchers often have to decide how to allocate budget across number of samples (including replicates and control data) and sequencing depth per sample. While existing guidelines for sequencing depth have focused on taxonomy estimation, MAG reconstruction, and gene detection [13, 14, 22, 24, 31, 37], our proposed modeling approach enables the principled study of the design of shotgun sequencing experiments to maximize power to detect differences in gene presence across sample groups.

Our latent variable model has possible utility for modeling the presence of amplicon sequence variants, and could offer a method for studying patterns of sequence variant presence when shotgun sequencing is infeasible or not preferred. For example, if a sequence variant is observed *W_i_* times in sample *i*, then it would be reasonable to model *Y_i_* = **1**_{*W_i_*>0}_. This would permit inference on the equality of the probability that the sequence variant is absent in a sample across sample groups. Notably, by choosing an *ϵ* > 0 (e.g., via the use of negative control samples), `happi` can adjust for the impact of index switching in studies that leverage multiplexing [15, 17]. We leave the application of `happi` to modeling the presence of amplicon sequence variants to future research.

Collectively, we have shown that `happi` is accurate and robust, even when genome quality is correlated with gene presence predictors. As the recovery of metagenome-assembled genomes becomes increasingly common, statistical tools that account for errors in recovered genomes become increasingly necessary. By leveraging genome quality metrics, `happi` provides sensible and inter-pretable results in an analysis of metagenome-assembled genome data, improves statistical inference under simulation, and can run efficiently on a local machine. Finally, by distributing open-source software in `R` implementing our proposed estimation and inference methods, we hope that `happi` can be used widely in a variety of genomics research settings. `happi` is available as an open-source `R` package via https://github.com/statdivlab/happi under a BSD-3-Clause license.

## 4 Methods

### 4.1 Methods: Saccharibacteria MAGs

The Saccharibacteria MAGs used in Data Analysis: Saccharibacteria MAGs, were taken from publicly available data [28]. Specifically, data on genome quality metrics (i.e. mean coverage) of these Saccharibacteria MAGs were retrieved from supplementary materials https://doi.org/10.6084/m9.figshare.11634321 and information on the presence or absence of COG functions in each MAG was extracted from the Saccharibacteria pangenome contigs databases and profiles located at https://doi.org/10.6084/m9.figshare.12217811. Functional annotation of the genes was performed using NCBI’s Clusters of Orthologous Groups (COG) database [32]. Further details on sampling, assembly, binning, and refinement can be found in [28]. In our data analysis, we specified *t*_max_ = 1000, Δ = 0.01 and *ϵ* = 0. We set *ϵ* = 0 because these MAGs had undergone careful manual refinement to remove contamination from other genomes. We suggest the use of *ϵ* > 0 when binning is performed automatically and without additional manual refinement.

### 4.2 Methods: simulation studies

#### 4.2.1 Subsampling study of *E. coli* isolate DRR102664

To investigate the probability of detecting a gene that it is truly present (*Pr*(*Y_i_* = 1|λ_*i*_ = 1, *M_i_* = *m*)), we conducted a subsampling simulation study of an *E. coli* isolate genome taken from [1]. We selected *E. coli* isolate DRR102664 to perform our subsampling simulation and the `eaeA` gene (K12790) as our target gene of interest. In enteropathogenic Escherichia coli, the `eaeA` gene produces a 94-kDa outer membrane protein called intimin which has been shown to be necessary to produce the attaching-and-effacing lesion. For our subsampling study, we subsampled paired sequences 50 times from the DRR102664 genome at approximate coverages *m* = (2×, 3×,…, 24×, 25×). Coverages were estimated using the calculation 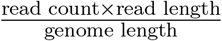. We annotated and identified the `eaeA` gene in each set of subsampled sequences and calculated the empirical probability of detection as the fraction of samples of coverage *m* that detected `eaeA`. The results of our subsampling investigation of the impact of coverage on the probability of detection given presence are shown in Figure 3.

#### 4.2.2 Evaluating estimators for *f*

Many different choices of functions *f* could be used to connect the probability of detecting a present gene to quality variables *M_i_*. We evaluated two options under simulation: 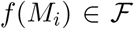 for 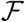 the class of bounded non-decreasing functions and 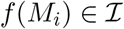 for 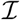 the class of bounded non-decreasing functions. As in Simulation Study, we set 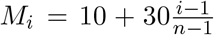, *X*_*i*1_ = 1, 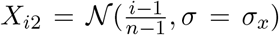, *β*_0_ = 0 and *ϵ* = 0. The true *f*(·) in this simulation is a generalized additive model with binomial link function [36] fit to the observations shown in Figure 3. This was done to select a true detection curve that well-reflects empirical probabilities of detecting a gene at a given coverage, such as gene `eaeA` in *E. coli* isolate genome DRR102664. We evaluated all estimators via mean squared error and median squared error for estimating *β*_1_. We investigated all combinations of *n* ∈ {30, 50, 100}, *β*_1_ ∈ {0.5, 1, 2} and *σ_x_* ∈ {0.25, 0.5}, and performed 250 draws for each combination. For 17 out of 18 combinations of *n*, *β*_1_ and *σ_x_*, we found that 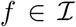 outperformed 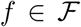 with respect to median squared error, with an average reduction in median squared error of 54%. For 18 out of 18 combinations, 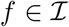 outperformed 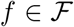 with respect to mean squared error, with an average reduction of 51%. For this reason, we chose to set 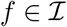 as the default option `happi`, and used this class of functions for both our data analyses and error rate simulations.

#### 4.2.3 Type 1 error and power simulations

For the Type 1 error rate and power simulations shown in Section 2.5, we performed 500 simulations for each combination of *σ_x_*, *β*_1_ and *n*. We set a minimum of 16 EM iterations, *t*_max_ = 50 and Δ = 0.1 for both the null and alternative models.

## Acknowledgements

The authors would like to thank Taylor Reiter and members of the StatDivLab for expert advice and constructive suggestions.

## Funding

This work was supported in part by the National Institute of General Medical Sciences (R35 GM133420); and the National Institute of Environmental Health Sciences (T32ES015459).

## Availability of data and materials

`happi` is available as an open-source `R` package at https://github.com/statdivlab/happi. The data supporting the conclusions of this article along with code for reproducing our results are made available at https://github.com/statdivlab/happi_supplementary.

## Notes

https://github.com/statdivlab/happi

https://github.com/statdivlab/happi_supplementary

### Competing Interest Statement

The authors have declared no competing interest.

## References

[1] Y. Arimizu, Y. Kirino, M. P. Sato, K. Uno, T. Sato, Y. Gotoh, F. Auvray, H. Brugere, E. Oswald, J. G. Mainil, K. S. Anklam, D. Döpfer, S. Yoshino, T. Ooka, Y. Tanizawa, Y. Nakamura, A. Iguchi, T. Morita-Ishihara, M. Ohnishi, K. Akashi, T. Hayashi, and Y. Ogura. Large-scale genome analysis of bovine commensal Escherichia coli reveals that bovine-adapted E. Coli lineages are serving as evolutionary sources of the emergence of human intestinal pathogenic strains. Genome Research, 29(9):1495–1505, 2019. doi: 10.1101/gr.249268.119.

[2] F. Beghini, L. J. McIver, A. Blanco-Míguez, L. Dubois, F. Asnicar, S. Maharjan, A. Mailyan, P. Manghi, M. Scholz, A. M. Thomas, M. Valles-Colomer, G. Weingart, Y. Zhang, M. Zolfo, C. Huttenhower, E. A. Franzosa, and N. Segata. Integrating taxonomic, functional, and strain-level profiling of diverse microbial communities with biobakery 3. eLife, 10:1–42, 2021. doi: 10.7554/eLife.65088.

[3] R. A. Blaustein, A. G. McFarland, S. Ben Maamar, A. Lopez, S. Castro-Wallace, and E. M. Hartmann. Pangenomic Approach To Understanding Microbial Adaptations within a Model Built Environment, the International Space Station, Relative to Human Hosts and Soil. mSystems, 4(1):1–16, 2019. doi: 10.1128/msystems.00281-18.

[4] O. Brynildsrud, J. Bohlin, L. Scheffer, and V. Eldholm. Rapid scoring of genes in microbial pan-genome-wide association studies with Scoary. Genome Biology, 17(1):1–9, 2016. doi: 10.1186/s13059-016-1108-8.

[5] L. X. Chen, K. Anantharaman, A. Shaiber, A. Murat Eren, and J. F. Banfield. Accurate and complete genomes from metagenomes. Genome Research, 30(3):315–333, 2020. doi: 10.1101/gr.258640.119.

[6] C. Collins and X. Didelot. A phylogenetic method to perform genome-wide association studies in microbes that accounts for population structure and recombination. PLOS Computational Biology, 14(2):1–21, 02 2018. doi: 10.1371/journal.pcbi.1005958. URL https://doi.org/10.1371/journal.pcbi.1005958.

[7] J. de Leeuw, K. Hornik, and P. Mair. Isotone optimization in R: Pool-adjacent-violators algorithm (pava) and active set methods. Journal of Statistical Software, 32(5):1–24, 2009. URL http://www.jstatsoft.org/v32/i05/.

[8] T. O. Delmont and A. M. Eren. Linking pangenomes and metagenomes: the prochlorococcus metapangenome. PeerJ, 6:e4320–e4320, 01 2018. doi: 10.7717/peerj.4320. URL https://pubmed.ncbi.nlm.nih.gov/29423345.

[9] A. P. Dempster, N. M. Laird, and D. B. Rubin. Maximum Likelihood from Incomplete Data Via the EM Algorithm. Journal of the Royal Statistical Society: Series B (Methodological), 39(1):1–22, 1977. doi: 10.1111/j.2517-6161.1977.tb01600.x.

[10] C. M. Duarte, D. K. Ngugi, I. Alam, J. Pearman, A. Kamau, V. M. Eguiluz, T. Gojobori, S. G. Acinas, J. M. Gasol, V. Bajic, and X. Irigoien. Sequencing effort dictates gene discovery in marine microbial metagenomes. Environmental Microbiology, 00:1–15, 2020. doi: 10.1111/1462-2920.15182.

[11] S. G. Earle, C. H. Wu, J. Charlesworth, N. Stoesser, N. C. Gordon, T. M. Walker, C. C. Spencer, Z. Iqbal, D. A. Clifton, K. L. Hopkins, N. Woodford, E. G. Smith, N. Ismail, M. J. Llewelyn, T. E. Peto, D. W. Crook, G. McVean, A. S. Walker, and D. J. Wilson. Identifying lineage effects when controlling for population structure improves power in bacterial association studies. Nature Microbiology, 1(5):1–8, 2016. doi: 10.1038/nmicrobiol.2016.41.

[12] A. M. Eren, Ö. C. Esen, C. Quince, J. H. Vineis, H. G. Morrison, M. L. Sogin, and T. O. Delmont. Anvi’o: an advanced analysis and visualization platform for ‘omics data. PeerJ, 3: e1319, 2015. doi: 10.7717/peerj.1319.

[13] H. S. Gweon, L. P. Shaw, J. Swann, N. De Maio, M. Abuoun, R. Niehus, A. T. M. Hubbard, M. J. Bowes, M. J. Bailey, T. E. A. Peto, S. J. Hoosdally, A. S. Walker, R. P. Sebra, D. W. Crook, M. F. Anjum, D. S. Read, N. Stoesser, M. Abuoun, M. Anjum, M. J. Bailey, L. Barker, H. Brett, M. J. Bowes, K. Chau, D. W. Crook, N. De Maio, D. Gilson, H. S. Gweon, A. T. M. Hubbard, S. Hoosdally, J. Kavanagh, H. Jones, D. S. Read, R. Sebra, L. P. Shaw, A. E. Sheppard, R. Smith, E. Stubberfield, J. Swann, A. S. Walker, and N. Woodford. The impact of sequencing depth on the inferred taxonomic composition and AMR gene content of metagenomic samples. Environmental Microbiomes, 14(1):1–15, 2019. doi: 10.1186/s40793-019-0347-1.

[14] B. Hillmann, G. A. Al-Ghalith, R. R. Shields-Cutler, Q. Zhu, D. M. Gohl, K. B. Beckman, R. Knight, D. Knights, and J. F. Rawls. Evaluating the information content of shallow shotgun metagenomics. mSystems, 3(6):e00069–18, 2018. doi: 10.1128/mSystems.00069-18. URL https://journals.asm.org/doi/abs/10.1128/mSystems.00069-18.

[15] Illumina. Effects of index misassignment on multiplexing and downstream analysis. Technical Report 770-2017-004-D, 2018. URL https://www.illumina.com/content/dam/illumina-marketing/documents/products/whitepapers/index-hopping-white-paper-770-2017-004.pdf.

[16] F. Imperi, L. C. Antunes, J. Blom, L. Villa, M. Iacono, P. Visca, and A. Carattoli. The genomics of Acinetobacter baumannii: Insights into genome plasticity, antimicrobial resistance and pathogenicity. IUBMB Life, 63(12):1068–1074, 2011. doi: 10.1002/iub.531.

[17] A. J. Larsson, G. Stanley, R. Sinha, I. L. Weissman, and R. Sandberg. Computational correction of index switching in multiplexed sequencing libraries. Nature Methods, 15(5):305–307, 2018. doi: 10.1038/nmeth.4666.

[18] J. A. Lees, M. Vehkala, N. Välimäki, S. R. Harris, C. Chewapreecha, N. J. Croucher, P. Marttinen, M. R. Davies, A. C. Steer, S. Y. C. Tong, A. Honkela, J. Parkhill, S. D. Bentley, and J. Corander. Sequence element enrichment analysis to determine the genetic basis of bacterial phenotypes. Nature Communications, 7(1):12797, 2016. doi: 10.1038/ncomms12797. URL https://doi.org/10.1038/ncomms12797.

[19] M. J. Pallen and B. W. Wren. Bacterial pathogenomics. Nature, 449(7164):835–842, 2007. doi: 10.1038/nature06248.

[20] E. Pasolli, F. Asnicar, S. Manara, M. Zolfo, N. Karcher, F. Armanini, F. Beghini, P. Manghi, A. Tett, P. Ghensi, M. C. Collado, B. L. Rice, C. DuLong, X. C. Morgan, C. D. Golden, C. Quince, C. Huttenhower, and N. Segata. Extensive Unexplored Human Microbiome Diversity Revealed by Over 150,000 Genomes from Metagenomes Spanning Age, Geography, and Lifestyle. Cell, 176(3):649–662.e20, 2019. doi: 10.1016/j.cell.2019.01.001. URL https://doi.org/10.1016/j.cell.2019.01.001.

[21] R. A. Power, J. Parkhill, and T. de Oliveira. Microbial genome-wide association studies: lessons from human gwas. Nature Reviews Genetics, 18(1):41–50, 2017. doi: 10.1038/nrg.2016.132. URL https://doi.org/10.1038/nrg.2016.132.

[22] C. Quince, A. W. Walker, J. T. Simpson, N. J. Loman, and N. Segata. Shotgun metagenomics, from sampling to analysis. Nature Biotechnology, 35(9):833–844, 2017. doi: 10.1038/nbt.3935. URL https://doi.org/10.1038/nbt.3935.

[23] L. Rouli, V. Merhej, P. E. Fournier, and D. Raoult. The bacterial pangenome as a new tool for analysing pathogenic bacteria. New Microbes and New Infections, 7:72–85, 2015. doi: 10.1016/j.nmni.2015.06.005. URL http://dx.doi.org/10.1016/j.nmni.2015.06.005.

[24] T. M. Royalty, A. D. Steen, and J. K. Jansson. Theoretical and simulation-based investigation of the relationship between sequencing effort, microbial community richness, and diversity in binning metagenome-assembled genomes. mSystems, 4(5):e00384–19, 2019. doi: 10.1128/mSystems.00384-19. URL https://journals.asm.org/doi/abs/10.1128/mSystems.00384-19.

[25] J. E. San, S. Baichoo, A. Kanzi, Y. Moosa, R. Lessells, V. Fonseca, J. Mogaka, R. Power, and T. de Oliveira. Current affairs of microbial genome-wide association studies: Approaches, bottlenecks and analytical pitfalls. Frontiers in Microbiology, 10, 2020. doi: 10.3389/fmicb.2019.03119. URL https://www.frontiersin.org/article/10.3389/fmicb.2019.03119.

[26] M. Scholz, D. V. Ward, E. Pasolli, T. Tolio, M. Zolfo, F. Asnicar, D. T. Truong, A. Tett, A. L. Morrow, and N. Segata. Strain-level microbial epidemiology and population genomics from shotgun metagenomics. Nature Methods, 13(5):435–438, 2016. doi: 10.1038/nmeth.3802. URL https://doi.org/10.1038/nmeth.3802.

[27] C. E. Sexton, H. Z. Smith, P. D. Newell, A. E. Douglas, and J. M. Chaston. MAGNAMWAR: an R package for genome-wide association studies of bacterial orthologs. Bioinformatics, 34(11):1951–1952, 01 2018. doi: 10.1093/bioinformatics/bty001. URL https://doi.org/10.1093/bioinformatics/bty001.

[28] A. Shaiber, A. D. Willis, T. O. Delmont, S. Roux, L.-X. Chen, A. C. Schmid, M. Yousef, A. R. Watson, K. Lolans, O. C. Esen, S. T. M. Lee, N. Downey, H. G. Morrison, F. E. Dewhirst, J. L. Mark Welch, and A. M. Eren. Functional and genetic markers of niche partitioning among enigmatic members of the human oral microbiome. Genome Biology, 21(1):292, 2020. doi: 10.1186/s13059-020-02195-w. URL https://doi.org/10.1186/s13059-020-02195-w.

[29] T. J. Sharpton. An introduction to the analysis of shotgun metagenomic data. Frontiers in Plant Science, 5, 2014. doi: 10.3389/fpls.2014.00209. URL https://www.frontiersin.org/article/10.3389/fpls.2014.00209.

[30] R. M. Sherman and S. L. Salzberg. Pan-genomics in the human genome era. Nature Reviews Genetics, 21(4):243–254, 2020. doi: 10.1038/s41576-020-0210-7. URL http://dx.doi.org/10.1038/s41576-020-0210-7.

[31] D. Sims, I. Sudbery, N. E. Ilott, A. Heger, and C. P. Ponting. Sequencing depth and coverage: key considerations in genomic analyses. Nature Reviews Genetics, 15(2):121–132, 2014. doi: 10.1038/nrg3642. URL https://doi.org/10.1038/nrg3642.

[32] R. L. Tatusov, N. D. Fedorova, J. D. Jackson, A. R. Jacobs, B. Kiryutin, E. V. Koonin, D. M. Krylov, R. Mazumder, S. L. Mekhedov, A. N. Nikolskaya, B. S. Rao, S. Smirnov, A. V. Sverdlov, S. Vasudevan, Y. I. Wolf, J. J. Yin, and D. A. Natale. The COG database: an updated version includes eukaryotes. BMC Bioinformatics, 4(1):41, 2003. doi: 10.1186/1471-2105-4-41. URL https://doi.org/10.1186/1471-2105-4-41.

[33] H. Tettelin, V. Masignani, M. J. Cieslewicz, C. Donati, D. Medini, N. L. Ward, S. V. Angiuoli, J. Crabtree, A. L. Jones, A. S. Durkin, R. T. DeBoy, T. M. Davidsen, M. Mora, M. Scarselli, I. M. y Ros, J. D. Peterson, C. R. Hauser, J. P. Sundaram, W. C. Nelson, R. Madupu, L. M. Brinkac, R. J. Dodson, M. J. Rosovitz, S. A. Sullivan, S. C. Daugherty, D. H. Haft, J. Selengut, M. L. Gwinn, L. Zhou, N. Zafar, H. Khouri, D. Radune, G. Dimitrov, K. Watkins, K. J. B. O’Connor, S. Smith, T. R. Utterback, O. White, C. E. Rubens, G. Grandi, L. C. Madoff, D. L. Kasper, J. L. Telford, M. R. Wessels, R. Rappuoli, and C. M. Fraser. Genome analysis of multiple pathogenic isolates of Streptococcus agalactiae: Implications for the microbial “pan-genome”. Proceedings of the National Academy of Sciences, 102(39):13950–13955, 2005. doi: 10.1073/pnas.0506758102. URL https://www.pnas.org/doi/abs/10.1073/pnas.0506758102.

[34] T. Van Rossum, P. Ferretti, O. M. Maistrenko, and P. Bork. Diversity within species: interpreting strains in microbiomes. Nature Reviews Microbiology, 18(9):491–506, 2020. doi: 10.1038/s41579-020-0368-1. URL https://doi.org/10.1038/s41579-020-0368-1.

[35] W. Wang and J. Yan. splines2: Regression Spline Functions and Classes, 2021. URL https://CRAN.R-project.org/package=splines2. R package version 0.4.5.

[36] S. N. Wood. Fast stable restricted maximum likelihood and marginal likelihood estimation of semiparametric generalized linear models. Journal of the Royal Statistical Society (B), 73(1):3–36, 2011.

[37] R. Zaheer, N. Noyes, R. O. Polo, S. R. Cook, E. Marinier, G. Van Domselaar, K. E. Belk, P. S. Morley, T. A. McAllister, R. Ortega Polo, S. R. Cook, E. Marinier, G. Van Domselaar, K. E. Belk, P. S. Morley, T. A. McAllister, R. O. Polo, S. R. Cook, E. Marinier, G. Van Domselaar, K. E. Belk, P. S. Morley, and T. A. McAllister. Impact of sequencing depth on the characterization of the microbiome and resistome. Scientific Reports, 8(1):1–11, 2018. doi: 10.1038/s41598-018-24280-8. URL http://dx.doi.org/10.1038/s41598-018-24280-8.

